# Uncovering mitochondrial diversity and candidate taxonomic lineages of *Hypostomus* (Siluriformes: Loricariidae) in Colombia

**DOI:** 10.64898/2026.11.26.769580

**Authors:** Liseth Paola Puello-Caballero, Paula Gudiela Orrego-Suárez, Armando Ortega-Lara, Carlos DoNascimiento, Luz Eneida Ochoa, Edna Judith Márquez

## Abstract

Molecular data for Colombian *Hypostomus* remain sparse, limiting assessments of species boundaries. We combined two mitochondrial markers (*cox1* and D-loop) with morphological, morphometric, geographic, and taxonomic evidence to evaluate diversity among Colombian populations. Using ASAP, bPTP, and GMYC, *cox1* recovered 16–17 Molecular Operational Taxonomic Units (MOTUs), whereas D-loop yielded 32–37, indicating finer mitochondrial lineage partitioning in D-loop that was not consistently mirrored by phenotypic differentiation. Because both markers belong to the same non-independent mitochondrial linkage group, congruence reflects partition stability rather than independent multi-locus support. High taxonomic-assignment support was obtained for *H. holostictus*, *H. hondae*, *H. niceforoi*, *H. plecostomoides*, *H. oculeus*, and *H. weberi*, whereas *H. carinatus*, *H. hemicochliodon*, and *H. sculpodon* received moderate support and *H. wilsoni* remained provisional because voucher-linked verification of the sequenced individuals was unavailable. Seven candidate lineages remain unresolved (G1, G2, and M3–M7); M8 was not counted among them because the examined specimens were morphologically compatible with *H. hemicochliodon*. Among these candidates, *Hypostomus* sp. G1 and G2 showed method-dependent mitochondrial partitioning and differences in selected morphometric characters relative to examined topotypic *H. robinii.* G2 also differed in pigmentation from examined *H. rhantos*, despite morphometric overlap. However, morphometric classification performance in Magdalena specimens and unresolved diagnostic boundaries do not provide sufficient evidence for recognizing G1 and G2 as distinct species. Mitochondrial delimitation patterns can reveal lineage partitioning and help identify taxonomic priorities, but should not be equated with species boundaries. Broader sampling, verified vouchers, and independent nuclear or genomic data are needed to evaluate these taxonomic hypotheses.

## Introduction

*Hypostomus* Lacépède, 1803 is one of the most diverse and taxonomically complex genera within Loricariidae, with a broad distribution throughout the Neotropical region [1,2]. Its identification and delimitation are hindered by high morphological and color-pattern variation, as well as by character overlap among related taxa [1,3,4]. Consequently, uncertainties persist regarding the correspondence between some nominal species and the lineages recovered through phylogenetic and delimitation analyses [2].

Molecular data have helped clarify these discrepancies by showing that genetic differentiation does not necessarily coincide with taxonomic boundaries inferred from morphology. Pronounced mitochondrial structuring has been documented among populations of *H. ancistroides* [5], while multilocus analyses of morphologically recognized sister species have revealed a mean cox1 divergence of 1.3 ± 1.2%, with only 11% of pairwise comparisons reaching or exceeding 2% [2]. Together these findings show that fixed thresholds of genetic divergence cannot be applied as universal criteria for species delimitation within *Hypostomus*.

Accordingly, groupings recovered from a single locus should be regarded as operational hypotheses of diversity rather than as delimited species. ASAP, bPTP, and GMYC use different analytical criteria (genetic distances or properties of genealogical branching) and may recover different partitions from the same molecular source [6–8]. Congruence among these methods indicates analytical stability of a partition but does not constitute biologically independent evidence of species boundaries. Their taxonomic interpretation therefore requires comparison with complementary sources of evidence, such as morphology, geographic distribution, ecology, and independent genetic data [9–11].

This caution is particularly important when inference is based exclusively on mitochondrial DNA. Different regions of the mitochondrial genome essentially share the same genealogical history and are not equivalent to independent nuclear loci. Although the analysis of multiple mitochondrial regions may increase the resolution of genetic structure, it does not eliminate the limitations associated with processes such as introgression, incomplete lineage sorting, or population structure [11,12]. Consequently, the detection of mitochondrial differentiation or lineage partitioning does not by itself demonstrate evolutionary independence at the species level. This distinction between genetic structure and species boundaries has been widely recognized in the molecular species-delimitation literature [11,13].

In Colombia, evaluation of this problem is limited by the still incomplete taxonomic and geographic representation in the major molecular analyses of *Hypostomus*. Although the national checklist recognizes considerable diversity within the genus [14], several Colombian species have not been incorporated into the most taxonomically comprehensive phylogenies, whereas others are represented by few individuals or localities [2]. This uneven coverage hinders assessment of the correspondence among nominal species, genetic differentiation, and morphological variation, as well as the identification of entities requiring taxonomic revision.

In this context, the present study had two objectives: (1) to characterize the mitochondrial diversity and its geographic distribution in Colombian *Hypostomus* using *cox1* and the D-loop control region, and to compare the mitochondrial lineage partitions inferred by ASAP, bPTP, and GMYC; and (2) to evaluate the taxonomic relevance of these partitions using morphological, morphometric, and geographic evidence, complemented with taxonomic material and reference sequences.

Molecularly recovered units were treated as Molecular Operational Taxonomic Units (MOTUs), representing operational hypotheses of mitochondrial lineage partitioning subject to subsequent evaluation. Their taxonomic relevance was assessed using phenotypic evidence and taxonomic comparisons, without assuming a one-to-one correspondence between MOTUs and species. Geographic distribution provided context for interpreting these patterns, but geographic separation alone was not considered evidence of taxonomic identity or species-level independence. Consistent with an integrative taxonomic approach [9–11], this framework distinguishes three levels of inference: mitochondrial diversity, mitochondrial lineage partitioning, and integrative taxonomic interpretation, including the evaluation of assignments to nominal species and hypotheses for unidentified morphotypes.

## MATERIAL AND METHODS

### Sampling

We analyzed 90 fin or muscle tissue samples from *Hypostomus* specimens collected across different hydrographic basins in Colombia (Fig 1; Table 1). Tissue samples were associated with specimens deposited in the CZUT-IC, CIACOL, CIUA, and MPUJ collections.

**Fig. 1.**
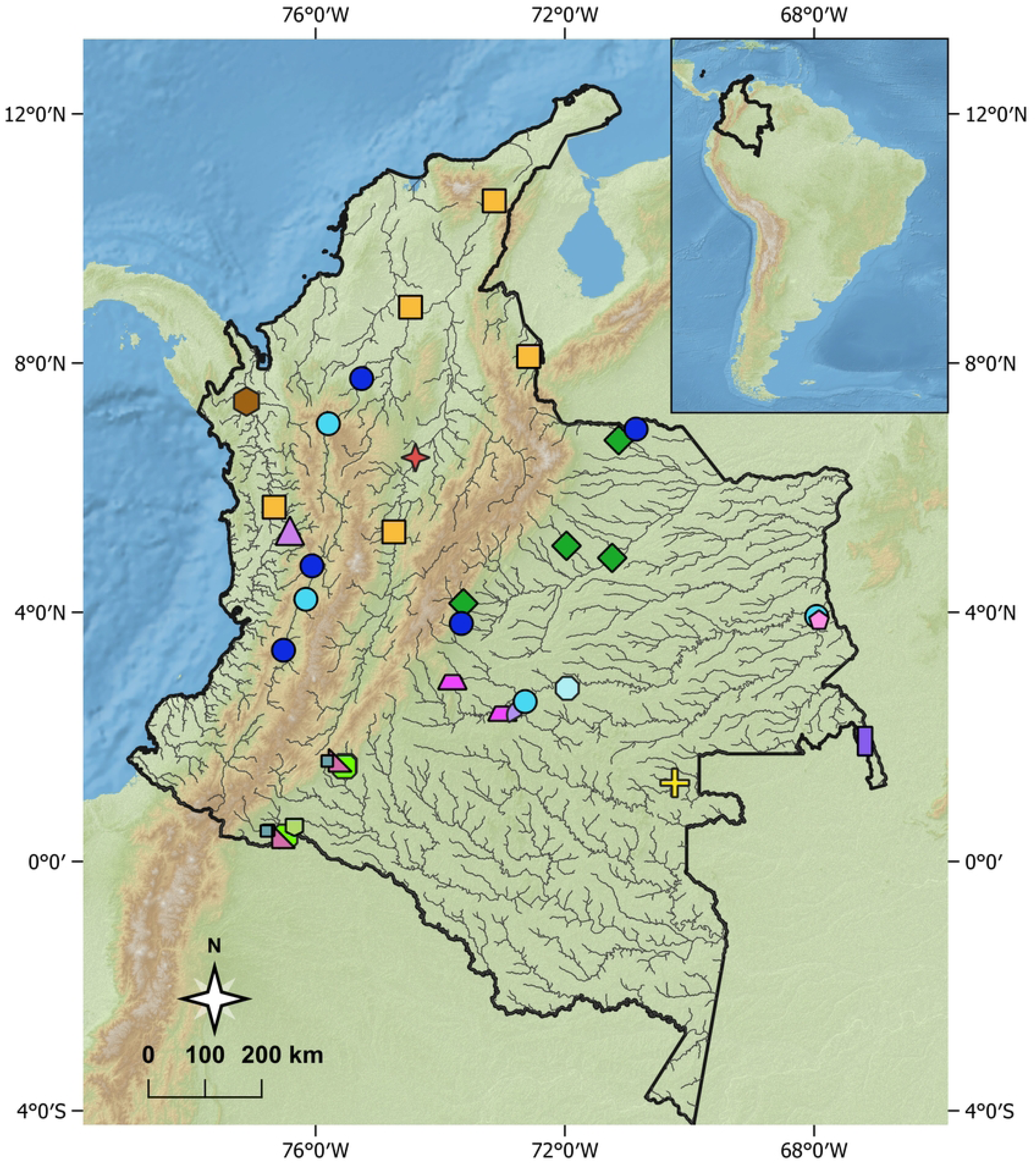
Geographic distribution of the *Hypostomus* lineages and morphotypes identified in this study in Colombia. Symbol colors and shapes represent: *H. hemicochliodon* (grayish-teal square), *H. hondae* (orange square), *H. carinatus* (pink pentagon), *Hypostomus* sp. G2 (light blue circle), *H. holostictus* (lilac triangle), *H. wilsoni* (dark brown hexagon), *Hypostomus* sp. G1 (blue circle), *H. weberi* (purple rectangle), *Hypostomus* sp. M6 Magdalena (red star), *Hypostomus* sp. M5 (light green pentagon), *Hypostomus* sp. M8 Putumayo (grayish-teal square), *Hypostomus* sp. M7 Vaupés (yellow cross), *Hypostomus* sp. M4 Guaviare (lilac irregular polygon), *Hypostomus* sp. M3 Guaviare (dark magenta trapezoid), *Hypostomus* sp. M3 Meta (light magenta trapezoid), *H. sculpodon* (light teal polygon), *H. plecostomoides* (green diamond), *H. oculeus* (pinkish-red triangle), and *H. niceforoi* (light green rounded square). Occurrence points correspond to georeferenced localities associated with specimens examined in this study. The Colombia boundary was obtained from the DANE Marco Geoestadístico Nacional 2024 (CC BY 4.0), available at https://geoportal.dane.gov.co/servicios/descarga-y-metadatos/datos-geoestadisticos/; country boundaries in the inset were obtained from Natural Earth Admin 0 Countries, 1:10m scale (public domain), available at https://www.naturalearthdata.com/downloads/10m-cultural-vectors/10m-admin-0-countries/; rivers and highlighted river segments were obtained from RiverATLAS v1.0/HydroATLAS (CC BY 4.0), available at https://www.hydrosheds.org/hydroatlas; elevation data were obtained from the USGS Global Multi-resolution Terrain Elevation Data 2010 (GMTED2010) mean-elevation grid at 7.5 arc-second resolution (∼250 m; public domain), with hillshade derived from these data in QGIS; and the physical background was obtained from Natural Earth II high-resolution raster data, 1:10m scale (public domain), available at https://www.naturalearthdata.com/downloads/10m-raster-data/10m-natural-earth-2/. The map was prepared in QGIS without proprietary satellite imagery or proprietary basemaps.

**Table 1.** Morphotypes and species of *Hypostomus* from Colombia included in the study.

| **Morphotype/species** | **S** | **Nc** | **Nd** | **River/locality of origin** | **Hydrographic region** | **GenBank accession(s), *cox1*** | **GenBank accession(s), D-loop** |
|---|---|---|---|---|---|---|---|
| *Hypostomus* sp. G1 | P | 25 | 25 | Cauca, Meléndez, Atrato, Dagua, Meta, and Arauca rivers | Magdalena, Pacific, Caribbean, Orinoco | QB064937–QB064961 | QB075155–QB075179 |
| *Hypostomus* sp. G2 | P | 15 | 16 | Cauca, Guaviare, and Inírida rivers | Magdalena, Orinoco | QB064962–QB064976 | QB075154,<br>QB075180–QB075194 |
| *Hypostomus* sp. M3 (5988, 6484) | C | 2 | 1 | Guaviare and Ariari rivers | Orinoco | QB064977–QB064978 | QB075195 |
| *Hypostomus* sp. M4 (6052) | C | 1 | — | Guaviare River drainage | Orinoco | QB064979 | — |
| *Hypostomus* sp. M5 (AOL1753) | C | 1 | — | Cohembí River | Amazon | QB064980 | — |
| *Hypostomus* sp. M6 (small spots) | U | 3 | 3 | Magdalena River, Puerto Berrío | Magdalena | QB064981–QB064983 | QB075196–QB075198 |
| *Hypostomus* sp. M7 (AOL1513) | C | — | 1 | Vaupés River | Amazon | — | QB075199 |
| *Hypostomus* sp. M8 (AOL1561, AOL1749, AOL1750) | C | 3 | — | Cohembí River | Amazon | QB064984–QB064986 | — |
| *Hypostomus carinatus* | C | 1 | 1 | Inírida River | Orinoco | QB064987 | QB075200 |
| *Hypostomus hemicochliodon* | C | 1 | — | Orteguaza River | Amazon | QB064988 | — |
| *Hypostomus holostictus* | U | 4 | 4 | San Juan River, Playa de Oro | Pacific | QB064989–QB064992 | QB075201–QB075204 |
| *Hypostomus hondae* | C | 12 | 11 | Magdalena River, Mesuno, Zulia River, Cesar River, Aguas Blancas, Catatumbo River, Atrato River, Badillo River, Tucuy River, Peralonso River, and Cauca River | Magdalena, Caribbean, Catatumbo | QB064993–QB065004 | QB075205–QB075215 |
| *Hypostomus niceforoi* | P | 4 | 4 | Orteguaza and Cohembí rivers | Amazon | QB065005–QB065008 | QB075216–QB075219 |
| *Hypostomus oculeus* | C | 3 | 2 | Huitoto and Bodoquerito rivers | Amazon | QB065009–QB065011 | QB075220–QB075221 |
| *Hypostomus plecostomoides* | C | 4 | 4 | Orinoco basin | Orinoco | QB065012–QB065015 | QB075222–QB075225 |
| *Hypostomus sculpodon* | C | 1 | — | Guaviare River | Orinoco | QB065016 | — |
| *Hypostomus weberi* | C | 2 | 4 | Guainía River, San Felipe | Amazon | QB065017–QB065018 | QB075226–QB075229 |
| *Hypostomus wilsoni* | U | 2 | — | Atrato River | Caribbean | QB065019–QB065020 | — |

S: *Hypostomus* supergroup sensu [2]; C: *Cochliodon*; P: *Plecostomus*; U: unassigned. Nc/Nd: number of *cox1*/D-loop sequences, respectively. G1–G2: broadly distributed morphotypes.

Taxonomic identification and morphological assessment were based on external morphology, color pattern, meristic characters, and morphometric measurements, compared to original descriptions, redescriptions, specialized taxonomic keys, and reference material [15–24]. When type material could not be examined directly, photographs of holotypes or other type specimens available from reference collections were used. These comparisons were used to assess correspondence between MOTUs and nominal species and to establish the taxonomic affinities of unassigned lineages. Species nomenclature and current taxonomic status follow [25].

Morphometric measurements were taken point-to-point with digital calipers to the nearest 0.01 mm, following established procedures for the group, with subsequent modifications [26,27]. Two sets of morphometric variables were recorded: a 15-variable matrix used for head analyses and an expanded matrix of 33 body and head variables. These included measurements of the head, mouth, predorsal region, trunk, and relative fin positions. Standard length (SL) was recorded in millimeters; all other measurements were expressed, according to the analytical matrix and corresponding anatomical region, as a percentage of SL (%SL) or head length (%HL). Raw morphometric measurements for the specimens included in the comparative datasets are provided in S2 Table. Morphological material examined is deposited in ANSP, CIACOL, CIUA, CZUT-IC, IAvH-P, ICN-MHN, MHNU, MLS, and MPUJ. Institutional abbreviations follow standard nomenclature [28,29].

### Molecular analyses and MOTU identification

Molecular sampling was restricted to taxa for which suitable tissue samples were available (Table 1), with detailed specimen information provided in S1 Table. Some nominal species recorded from Colombia, including *Hypostomus annectens*, *H. argus*, and *H. varimaculosus* [14], could not be included because suitable material was unavailable or because of access and sampling limitations associated with their known distributions.

Mitochondrial diversity of *Hypostomus* in Colombia was assessed using two mitochondrial regions, *cox1* and D-loop. Because of their linked inheritance, both markers were analyzed separately and treated as complementary sources of mitochondrial variation rather than as evolutionarily independent loci for species delimitation. Among these markers, *cox1* was selected because of its broad representation in reference databases, whereas D-loop was used to explore mitochondrial variation at a finer scale.

Genomic DNA was extracted from fin or muscle tissue preserved in 70% ethanol using the GF-1 Nucleic Acid Extraction Kit, following the manufacturer’s instructions. Partial fragments of *cox1* were amplified by polymerase chain reaction (PCR) using the primers VF2 and FishR2 [30], and the D-loop control region using DLA-III and HygDL-R [31].

PCRs were performed in a final volume of 25 µL, consisting of 22 µL of reaction mixture and 3 µL of template DNA (∼20 ng). Final concentrations were 1× PCR buffer, 1.5 mM MgCl₂, 0.2 mM of each dNTP, 0.2 µM of each forward and reverse primer, and 1 U of Platinum® Taq Polymerase (Invitrogen). Sterile nuclease-free water was added to reach the final volume. The thermal profile consisted of an initial denaturation at 95 °C for 5 min, followed by 30 cycles of 95 °C for 35 s, annealing for 35 s at 56 °C for cox1 or 55 °C for D-loop, and extension at 72 °C for 35 s, followed by a final extension at 72 °C for 5 min.

PCR products were verified by electrophoresis on 2% agarose gels, purified with alkaline phosphatase and exonuclease I according to the manufacturer’s instructions, and sequenced by the Sanger method using the BigDye™ Terminator v3 Cycle Sequencing Ready Reaction Kit (Applied Biosystems) on a four-capillary SeqStudio sequencer.

Of the 90 samples processed, 84 *cox1* and 76 D-loop sequences were successfully obtained. Sequences for both markers were recovered from 70 specimens, whereas 14 yielded only *cox1*, and six only for D-loop. The *cox1* dataset comprised 129 terminals, 85 sequences generated in this study (84 *Hypostomus* sequences and one *Pterygoplichthys pardalis*) and 44 sequences retrieved from GenBank, representing 35 *Hypostomus*, six other loricariid taxa, and three outgroup taxa. The D-loop dataset comprised 80 terminals, including 77 sequences generated in this study (76 *Hypostomus* sequences and one *P. pardalis*) and three reference sequences retrieved from GenBank, corresponding to *H. ancistroides*, *P. pardalis*, and *P. disjunctivus*. Detailed specimen and sequence information is provided in S1 Table.

Complementary GenBank (NCBI) sequences were selected using standardized curation criteria: ≥ 80% coverage of the amplicon length obtained in this study, < 1% ambiguous nucleotide sites, and taxonomic and geographic traceability supported by peer-reviewed publications or voucher specimens. For *cox1*, reading-frame integrity and the absence of premature stop codons were additionally verified.

The final dataset comprised 129 *cox1* sequences aligned over 609 bp and 80 D-loop sequences aligned over 557 bp. *cox1* sequences were aligned using MUSCLE [32]. As a quality-control step, bidirectional chromatograms were manually inspected for overlapping peaks, and sequences were conceptually translated using the vertebrate mitochondrial genetic code. No premature stop codons or reading-frame disruptions were detected, reducing the likelihood of including nuclear mitochondrial pseudogenes (NUMTs; [33]).

Because of their higher variability and the potential presence of ambiguously aligned regions, D-loop sequences were aligned with MAFFT v7.407_1 [34] with the automatic strategy, a gap-opening penalty of 1.53, and a gap-extension offset of 0.123, without automatic sequence orientation adjustment. Ambiguously aligned regions associated with repeats or length variation were inspected and removed using the automatic trimming option in trimAl v1.4.1 [35]. In subsequent analyses, remaining gaps were treated as missing data and ambiguous nucleotide sites were coded according to IUPAC nomenclature.

The nucleotide substitution model for each marker was selected in jModelTest2 [36] using the Akaike Information Criterion (AIC; [37]). The selected models were TPM2uf+I+G for *cox1* and TrN+I+G for D-loop. Because of the parameterization options available in the programs used for phylogenetic inference, GTR+I+G was used as an approximation to these models in analyses associated with bPTP and GMYC.

Mitochondrial lineage partitioning was explored using three molecular delimitation methods: Assemble Species by Automatic Partitioning (ASAP; [8]), Bayesian Poisson Tree Processes (bPTP; [7]), and Generalized Mixed Yule–Coalescent (GMYC; [38]). The recovered units were termed Molecular Operational Taxonomic Units (MOTUs) and interpreted as hypotheses of mitochondrial clustering rather than independently delimited species. Congruence among methods was used to assess partition stability across different analytical approaches, whereas congruence between *cox1* and D-loop was interpreted as consistency of the mitochondrial pattern rather than independent multilocus corroboration. The taxonomic relevance of MOTUs was subsequently assessed through their congruence with geographic distribution and morphological and morphometric evidence.

ASAP was run independently for each marker using the corresponding alignments and Kimura two-parameter distances (K2P; [39]). For comparisons among methods, the partition with the lowest ASAP score was retained, without assuming *a priori* that it represented a definitive taxonomic delimitation.

For bPTP, consensus topologies obtained by Bayesian inference in MrBayes v3.2 [40] under the GTR+I+G model were used as input trees. Independent MCMC runs were performed with four chains, sampling trees at regular intervals. The D-loop analysis was extended to 6,370,000 generations and reached a final average standard deviation of split frequencies (ASDSF) of 0.003640, whereas the *cox1* analysis was extended to 3,355,000 generations and reached an ASDSF of 0.007173. Both values were below 0.01, supporting convergence of split frequencies among runs. The initial 25% of samples were discarded as burn-in, and sampling efficiency of continuous parameters was evaluated using effective sample sizes (ESS), with values > 200 considered adequate. The resulting consensus trees were subsequently used as input for bPTP, with 10,000 Markov chain Monte Carlo (MCMC) generations, the H1 method, sampling every 100 generations, and a burn-in of 10%. The random seed was set to 123. No outgroup taxa were specified in the configuration, and the outgroup removal option was disabled.

For GMYC, an ultrametric tree was estimated for each marker in BEAST v2.7.7. [41]. An uncorrelated lognormal relaxed molecular clock (UCLN) and a Birth–Death tree prior were initially explored. This parameterization showed convergence problems and low sampling efficiency, with ESS values < 100 for several parameters even after tens of millions of generations. A more parsimonious configuration was therefore adopted, using a strict molecular clock and a Yule (pure-birth) process as the tree prior. This configuration reduced model dimensionality and produced chains with ESS > 200 for the continuous parameters evaluated. The strict clock was used exclusively to obtain the ultrametric trees required by GMYC and not to estimate divergence times.

BEAST MCMC chains were run for 50 million generations, sampling every 50,000 generations. Convergence and sampling efficiency were assessed in Tracer v1.7.2 [42], with ESS values > 200 considered satisfactory. After discarding the first 25% of samples as burn-in, a maximum clade credibility (MCC) tree was obtained for each marker using TreeAnnotator v2.7.7. The single-threshold GMYC model was applied separately to the resulting ultrametric trees for *cox1* and D-loop using the splits 1.0-14/r31 package [43] in R [44]. For each marker, model fit was evaluated using a likelihood-ratio test comparing the GMYC model with the null model of a single coalescent process. The log-likelihoods of both models and the estimated threshold time were recorded.

## Morphometric and statistical analyses of morphotypes and species within the *Hypostomus plecostomus* supergroup

### Morphometric datasets and allometric correction

Morphometric analyses followed a hierarchical design. The initial dataset comprised 35 specimens representing *Hypostomus niceforoi* (n = 5), *H. rhantos* (n = 10), *Hypostomus* sp. G1 (n = 6), *Hypostomus* sp. G2 (n = 10), and topotypic *H. robinii* (n = 4). Subsequent analyses focused on G1, G2, and topotypic *H. robinii* (N = 20). All G1 and G2 specimens in these reference datasets originated from the Orinoco basin. An additional nine specimens from the Magdalena basin, independently assigned to G1 (n = 7) or G2 (n = 2) based on molecular evidence, were reserved exclusively for external validation of the three-group morphometric models.

For exploratory analyses, measurements were expressed as percentages of standard length (%SL) or head length (%HL), as appropriate, and natural-log transformed before PCA and LDA or PCA-LDA. Because ratio standardization may retain allometric size effects [45], residual size dependence was assessed using multivariate models incorporating group, log-transformed size, and their interaction, with significance evaluated using Pillai’s trace [46,47]. Persistent size effects restricted these analyses to exploratory interpretation.

Original measurements were allometrically corrected separately for each analytical dataset [48,49], using SL for body measurements and HL for head measurements. Homogeneity of allometric slopes among groups was assessed for each variable by comparing log-linear models with and without the group × log(size) interaction. Interaction P-values were Holm-adjusted across variables [50], and variables showing significant slope heterogeneity were excluded from common-slope correction. For the remaining variables, common within-group allometric exponents were estimated while accounting for group differences. In the focal G1–G2–*H. robinii* head dataset, the mean HL was calculated from all 20 specimens. Residual size effects were reassessed after correction as described above.

### Multivariate and univariate analyses

Analyses were conducted in R v4.4.2 [51], using the stats package for PCA [51], MASS for LDA [52], and ggplot2 for visualization [53]. Body and head measurements were analyzed separately. PCA-LDA was used when variable dimensionality exceeded the available within-group rank; otherwise, direct LDA was applied. PCA [54] was performed on centered and standardized measurements, retaining components explaining at least 95% of cumulative variance, subject to the available-rank constraint. All discriminant models used equal group priors.

In the broad five-group dataset (N = 35), all 33 body variables passed Holm-adjusted slope-homogeneity screening. Both exploratory ln(%SL) and allometrically corrected body matrices were analyzed using PCA-LDA because their dimensionality exceeded the maximum within-group rank (≤30). Direct LDA was applied to the exploratory head matrix (15 variables) and the corrected head matrix (14 retained variables). Residual size dependence was assessed on retained principal components using Pillai’s trace.

In the focal G1–G2–*H. robinii* dataset (N = 20), all 33 body and 14 head variables passed Holm-adjusted slope-homogeneity testing. The corrected body matrix was analyzed using PCA-LDA because its dimensionality exceeded the within-group rank (17). Twelve principal components, explaining 95.49% of cumulative variance, were retained under a maximum of 16 components. LDA used equal group priors and a tolerance of 1 × 10⁻¹⁰. Residual size dependence was assessed on components accounting for at least 95% of cumulative variance. The corrected head matrix was analyzed using direct LDA.

Classification performance of the broad corrected head matrix was assessed using leave-one-out cross-validation (LOOCV [55]), reporting overall accuracy, balanced accuracy, and Cohen’s κ. Because variable selection and allometric correction were performed before cross-validation, these results were interpreted as a sensitivity assessment rather than fully nested predictive validation.

Predictive performance of the focal body PCA-LDA and head LDA models was evaluated using fully nested LOOCV. Within each training fold, slope-homogeneity testing, Holm adjustment, variable selection, reference size, common allometric exponents, and allometric correction were recalculated using training specimens only. For body PCA-LDA, centering, scaling, PCA, and component selection were also repeated before fitting LDA. Held-out specimens were transformed using exclusively training-derived parameters. Models used equal group priors, and performance was summarized by overall accuracy, balanced accuracy, and group-specific recall. Full-data reclassification accuracy was considered descriptive only.

Univariate differences among G1, G2, and *H. robinii* were evaluated using the 14 corrected head variables. Residual normality and variance homogeneity were assessed using Shapiro–Wilk [56] and Brown–Forsythe [57] tests, respectively. Depending on these assumptions, one-way ANOVA with Tukey–Kramer comparisons [58,59], Welch’s ANOVA with Games–Howell comparisons [60,61], or Kruskal–Wallis with Dunn’s post hoc comparisons [62,63] was applied. Global-test P-values were Holm-adjusted across variables, and post hoc comparisons were interpreted only when the corresponding adjusted global test was significant.

### External validation

External transferability of the focal three-class head LDA model (G1, G2, and *H. robinii*; reference N = 20) was evaluated using nine Magdalena specimens molecularly assigned to G1 or G2. The model was based on the 14 retained allometrically corrected head variables. External specimens were excluded from allometric estimation, variable screening, model development, and internal cross-validation. Before prediction, their measurements were matched to the reference variables in identity and order and transformed using preprocessing parameters estimated exclusively from the 20-specimen focal reference dataset. Predictions retained all three possible classification categories, whereas molecular assignments to G1 or G2 were used only to evaluate classification performance after predictions had been obtained.

### Integrative taxonomic assignment

Taxonomic assignment support for nominal species and taxonomic hypothesis support for unidentified morphotypes were qualitatively categorized as high (H), moderate (M), or provisional (P), based primarily on diagnostic phenotypic evidence and taxonomic comparisons, complemented by sample representation and mitochondrial compatibility. For morphotypes, support reflected proposed identities, affinities, or differentiation from nominal species, considering corroborating evidence and unresolved alternatives. Operational criteria, decision rules, and case-specific justifications are provided in S4 Table.

Mitochondrial subdivisions warranted reconsideration of nominal species assignments only when associated with unresolved taxonomic inconsistencies. For morphotypes, they were evaluated against alternative taxonomic hypotheses without equating MOTUs with species. Mitochondrial subdivisions alone were considered insufficient for species delimitation, and agreement among delimitation algorithms or linked mitochondrial markers was not treated as independent corroboration. Geography provided interpretive context but did not determine support categories, which reflect confidence in integrative taxonomic interpretations rather than statistical significance, formal taxonomic categories, or species-delimitation decisions.

## Results

The delimitation methods recovered a considerably higher number of MOTUs from D-loop than from *cox1*: for *cox1*, ASAP and GMYC recognized 16 MOTUs and bPTP 17, whereas D-loop recovered 35, 32, and 37 MOTUs, respectively (Figs 2–5). For *cox1*, the null model log-likelihood was 928.99, the maximum log-likelihood of the GMYC model was 934.91, and the estimated threshold time was −0.007. For D-loop, the corresponding values were 574.00, 577.35, and −0.002, respectively.

**Fig 2.**
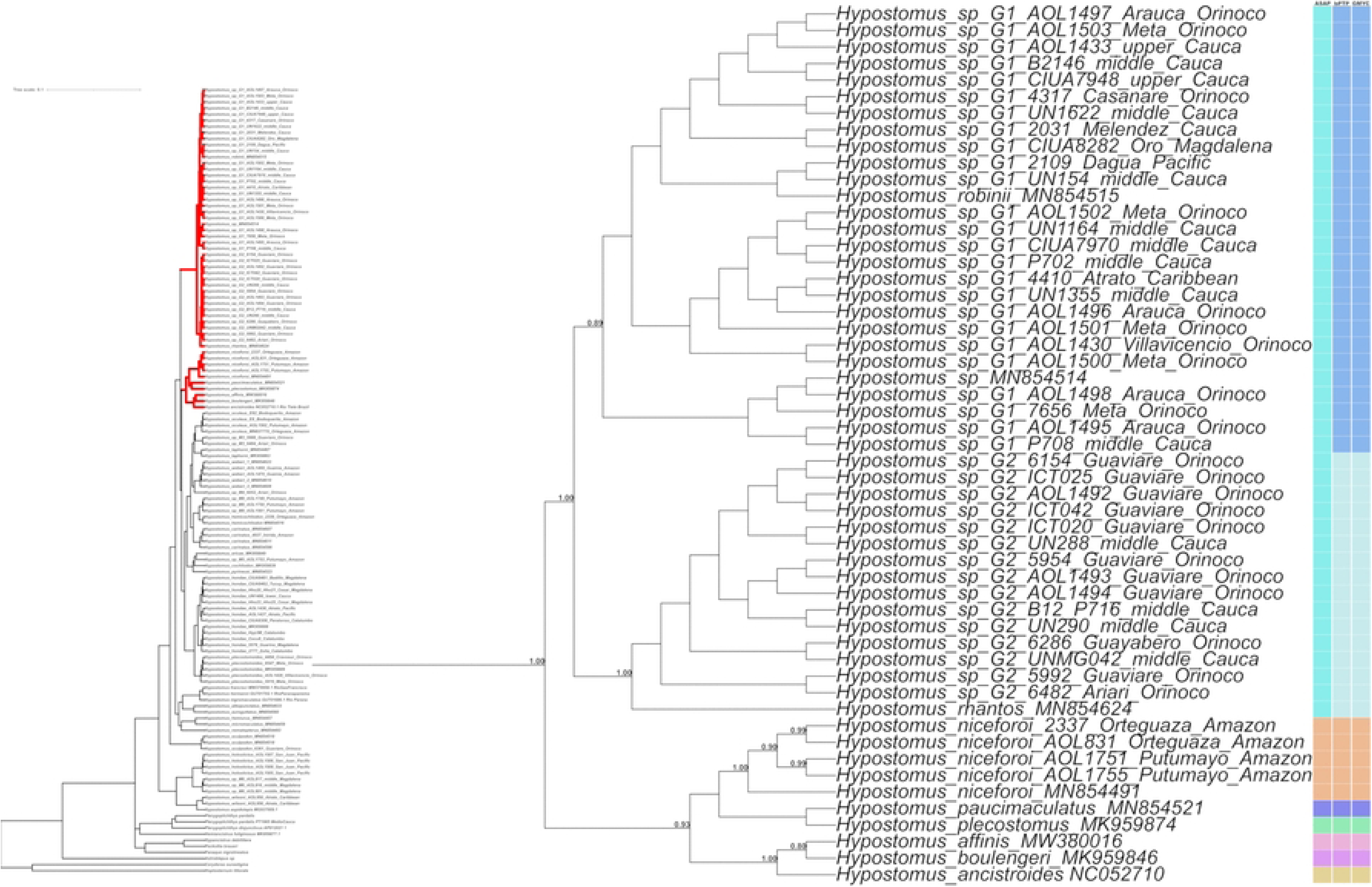
MOTU delimitation in the *Hypostomus plecostomus* supergroup based on *cox1*. The maximum clade credibility tree summarizes the Bayesian phylogenetic analysis. The columns alongside the tree show the partitions recovered by ASAP, bPTP, and GMYC. Within each column, different colors indicate different MOTUs.

**Fig 3.**
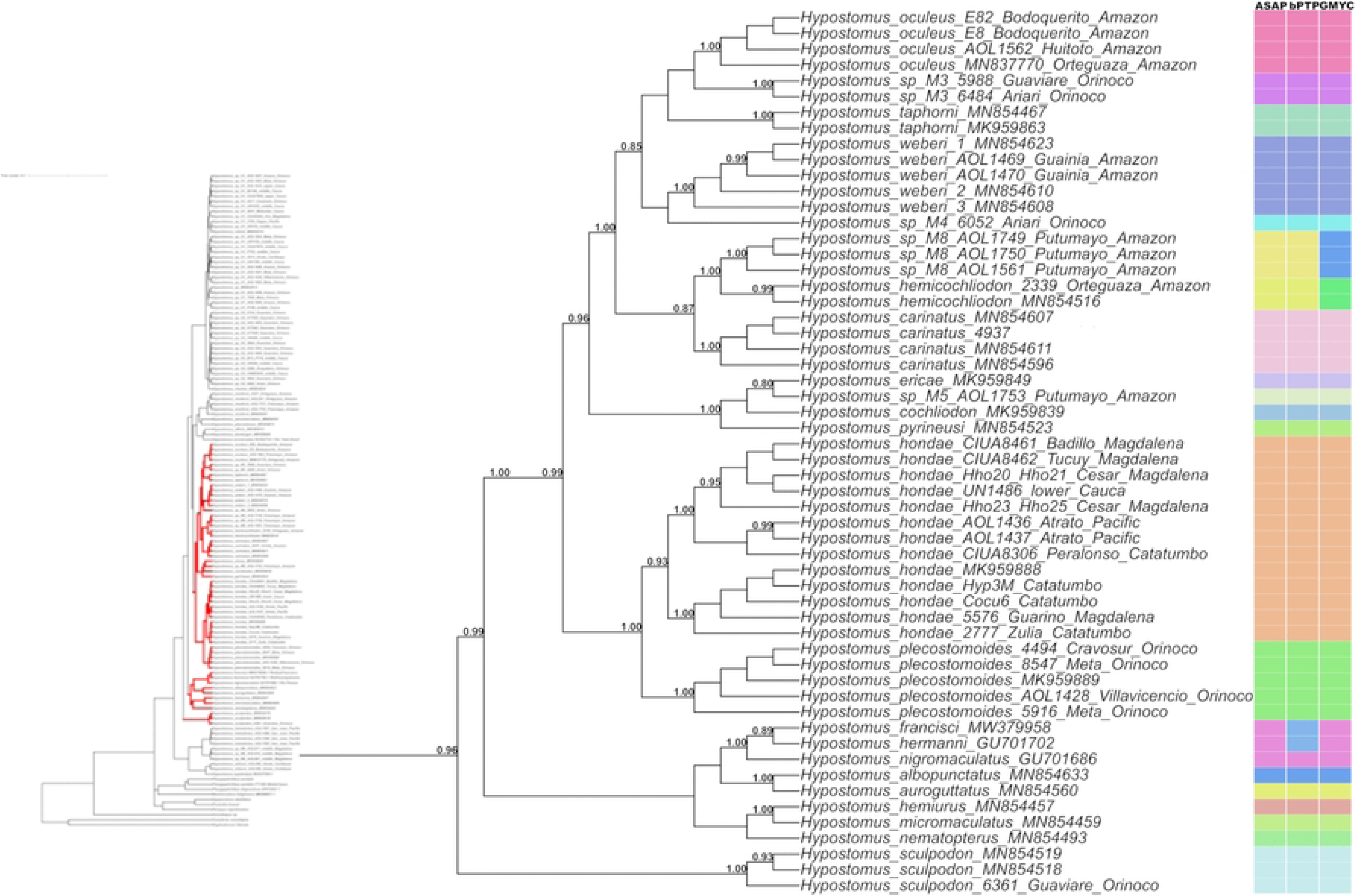
MOTU delimitation in the *Hypostomus cochliodon* supergroup based on *cox1*. The maximum clade credibility tree summarizes the Bayesian phylogenetic analysis. The columns alongside the tree show the partitions recovered by ASAP, bPTP, and GMYC. Within each column, different colors indicate different MOTUs.

**Fig 4.**
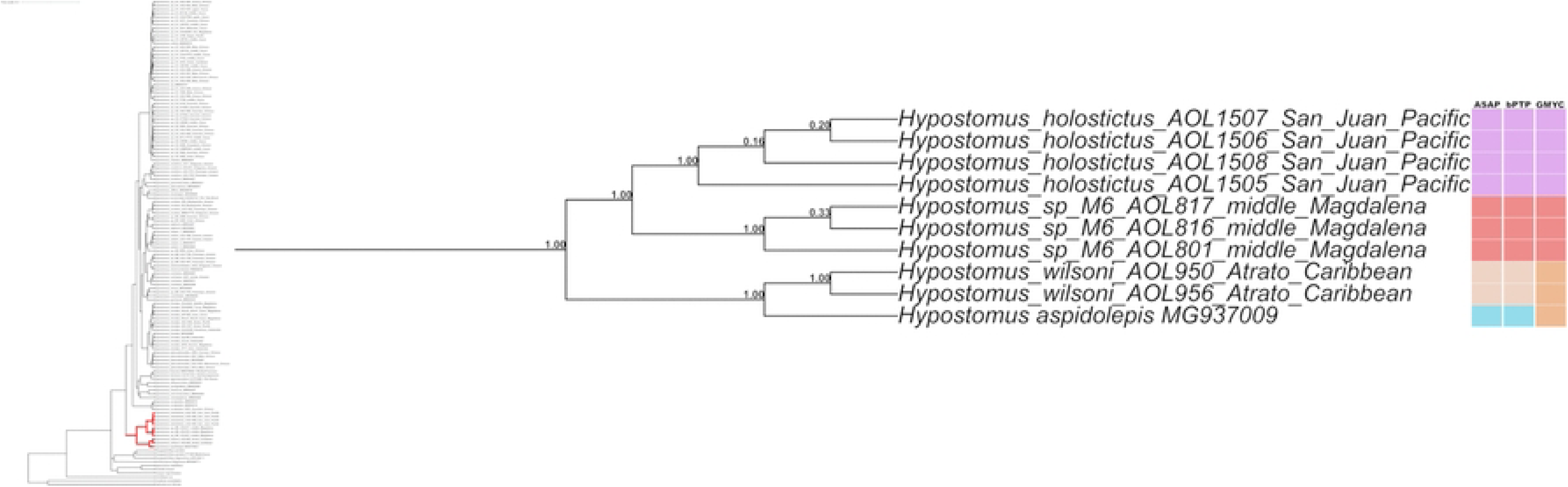
MOTU delimitation for *Hypostomus* specimens from the Pacific drainage and the middle Magdalena River basin based on *cox1*. In the Bayesian maximum clade credibility tree, these specimens fall outside all four previously proposed supergroups [2]. The columns alongside the tree show the partitions recovered by ASAP, bPTP, and GMYC. Within each column, different colors indicate different MOTUs.

**Fig 5.**
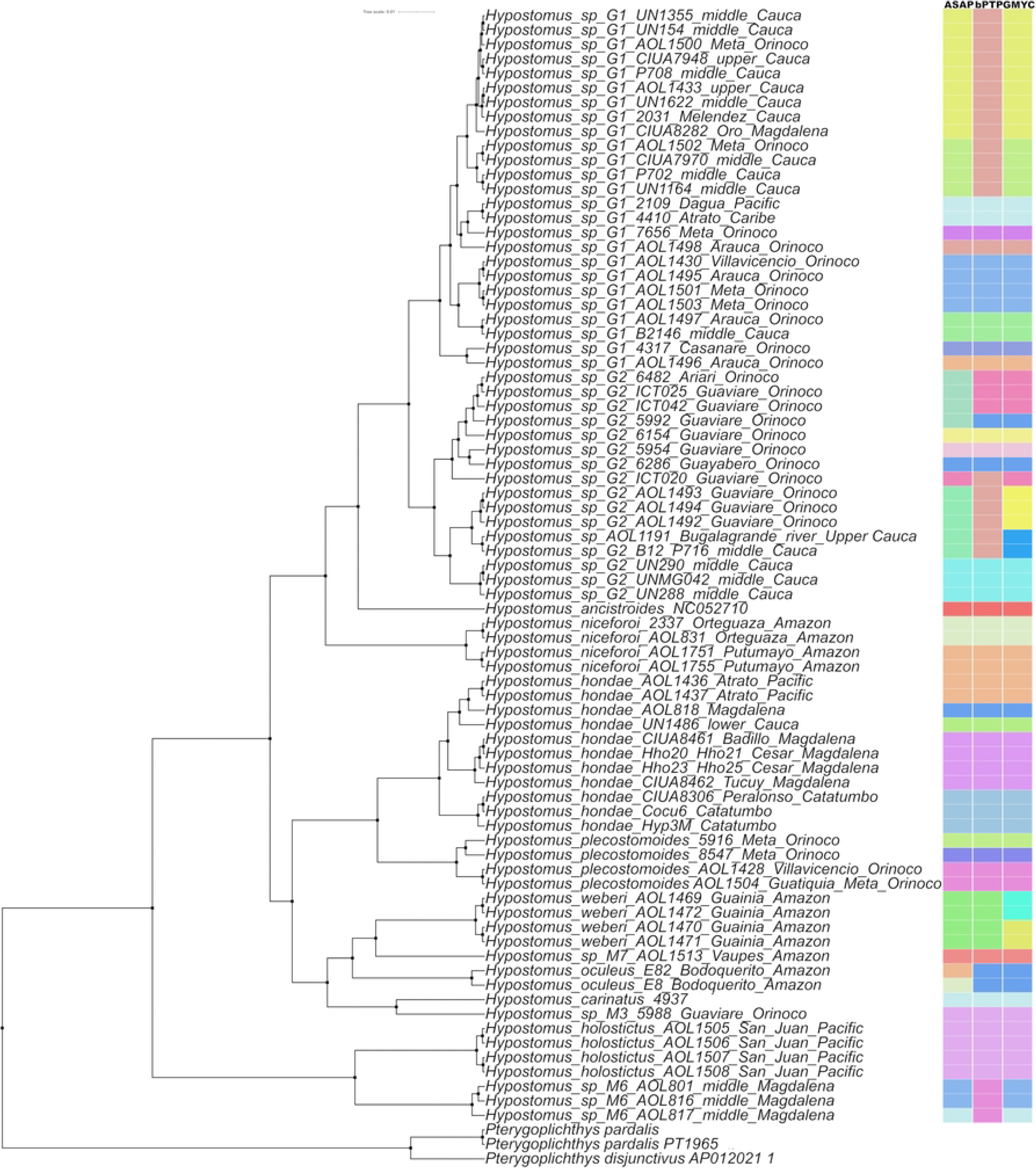
MOTU delimitation of *Hypostomus* based on D-loop. The maximum clade credibility tree summarizes the Bayesian phylogenetic analysis. The columns alongside the tree show the partitions recovered by ASAP, bPTP, and GMYC. Within each column, different colors indicate different MOTUs.

Most nominal species were consistently recovered as distinct MOTUs by the three delimitation methods. Among the morphotypes, different phylogenetic affinities were observed across hydrographic regions. Within the *Hypostomus plecostomus* supergroup (Fig 2), all haplotypes of *Hypostomus* sp. G1, distributed across the Magdalena, Pacific, Caribbean, and Orinoco regions, showed close affinity with *H. robinii*, whereas haplotypes of *Hypostomus* sp. G2 from the Magdalena and Orinoco regions were most closely associated with *H. rhantos*. The G1–*H. robinii* and G2–*H. rhantos* groups were themselves phylogenetically related.

Within the *Hypostomus cochliodon* supergroup (Fig 3), *Hypostomus* sp. M3 from the Orinoco region showed affinities with *H. oculeus* and *H. taphorni*, while *Hypostomus* sp. M4 from the Orinoco region and *Hypostomus* sp. M7 from the Amazon region were both associated with *H. weberi*. In the Amazon region, *Hypostomus* sp. M5 showed affinities with *H. cochliodon*, *H. ericae*, and *H. pyrineusi*, whereas *Hypostomus* sp. M8 showed affinity with *H. hemicochliodon*. In contrast, *Hypostomus* sp. M6 from the Magdalena region was associated with *H. holostictus* in a phylogenetic group not currently assigned to a recognized supergroup (Fig 4).

For *cox1*, all three methods concordantly recovered 12 MOTUs corresponding to *H. carinatus*, *H. holostictus*, *H. hondae*, *H. niceforoi*, *H. oculeus*, *H. weberi*, *H. plecostomoides*, *H. sculpodon*, as well as morphotypes of *Hypostomus* sp. M3, sp. M4, sp. M5, and sp. M6. Differences among methods were restricted to three groups: *H. aspidolepis*–*H. wilsoni* (Fig 4), *H. hemicochliodon*–*Hypostomus* sp. M8 (Fig 3), and the G1–*H. robinii* and G2–*H. rhantos* groups (Fig 2). Among these cases, ASAP and bPTP separated *H. wilsoni* from *H. aspidolepis* (Fig 4) and *H. hemicochliodon* from *Hypostomus* sp. M8, whereas GMYC grouped each pair into a single MOTU (Fig 3).

Within the G1–G2 complex, ASAP grouped both lineages into a single MOTU, whereas bPTP and GMYC recovered them as separate units. Reference sequences of *H. robinii* and *H. rhantos* fell within the same phylogenetic complex, although their correspondence with molecular partitions varied among methods (Fig 2).

D-loop showed greater and more heterogeneous subdivision into MOTUs among groups (Fig 5). *Hypostomus carinatus*, *H. holostictus*, *Hypostomus* sp. M3, and *Hypostomus* sp. M7 were each recovered as a single MOTU in the available sampling. In contrast, *H. hondae*, *H. niceforoi*, and *H. plecostomoides* were concordantly subdivided into five, two, and three MOTUs, respectively.

Other groups showed disagreement among delimitation methods: *H. weberi* was recovered as a single MOTU by ASAP and bPTP and as two by GMYC; *H. oculeus* as a single MOTU by bPTP and GMYC and as two by ASAP; and *Hypostomus* sp. M6 as a single MOTU by bPTP and as two by ASAP and GMYC.

The greatest subdivision was observed in G1 and G2. In G1, ASAP and GMYC recovered nine MOTUs and bPTP eight; in G2, ASAP and bPTP recovered seven MOTUs, although with different composition, whereas GMYC recovered nine. Overall, D-loop yielded considerably finer MOTU partitioning than *cox1*, particularly within the G1–G2 complex (Fig 5).

### Morphological results of the evaluated species and morphotypes

Phenotypic identifications of nominal species were supported by diagnostic characters documented in original descriptions, redescriptions, taxonomic keys, and, where available, comparisons with type or reference material. Direct examination of the holotypes corroborated the assignments of *Hypostomus niceforoi* (ANSP 70511) and *H. oculeus* (ANSP 70518). Specimens assigned to *H. holostictus* exhibited diagnostic dental and external characters, including unicuspid teeth, consistent with the original description and photographs of the holotype (BMNH 1913.10.1.57). For *H. wilsoni*, independently examined material was morphologically consistent with the species and was compared with photographs of the holotype (FMNH 58550), although the sequenced tissues lacked directly associated vouchers.

Several lineages not assigned to nominal species showed character combinations that differed from those of their closest phylogenetic relatives. *Hypostomus* sp. G1 and G2 resembled *H. robinii* in qualitative external morphological characters. However, morphometric comparisons with the examined topotypic specimens revealed significant differences in some measurements, as detailed below. *Hypostomus* sp. G2 differed from *H. rhantos* primarily in pigmentation pattern. Comparison with *H. rhantos* included the paratypes IAvH-P 13958 and ANSP 185240, together with the diagnostic information available for the species. *Hypostomus rhantos* is characterized by numerous, very small and densely distributed spots on the head and dorsal and lateral surfaces of the body, whereas the examined G2 specimens showed a qualitatively different pigmentation pattern, with fewer and larger spots. Because spot abundance in *H. rhantos* varies with body size, this comparison was treated qualitatively rather than as a fixed meristic diagnostic criterion*. Hypostomus* sp. M3 differed from *H. oculeus* in lacking conspicuous body spots, with pigmentation largely restricted to the ventral region, and from *H. taphorni* in lacking the characteristic bicolored caudal pattern described for that species. *Hypostomus sp.* M7 differed from *H. weberi* by having unicuspid teeth and by the pigmentation of the distal portions of the fins, which were more heavily pigmented. In contrast, *Hypostomus* sp. M4 differed mainly in body pigmentation, with smaller and more closely spaced spots than those of *H. weberi,* which has larger and more widely spaced spots, and by the absence of the diagnostic oral papilla described for *H. weberi*.

*Hypostomus* sp. M5 differed from *H. ericae* in lacking an oral papilla, from *H. cochliodon* in its uniform dark coloration and absence of conspicuous spots on the fins or abdomen, and from *H. pyrineusi* in having more than 10 opercular odontodes. *Hypostomus* sp. M6 differed from *H. holostictus* by having bicuspid teeth, whereas *H. holostictus* is characterized by unicuspid teeth. *Hypostomus* sp. M6 also showed a distinct morphological combination and is currently undergoing separate taxonomic study. In contrast, specimens of *Hypostomus* sp. M8 were morphologically congruent with *H. hemicochliodon*, and no diagnostic characters were observed that distinguished them from that species.

### Broad morphometric comparison of morphotypes and species within the *Hypostomus plecostomus* supergroup

In the broad five-group comparison, exploratory analyses of ln(%SL) body measurements retained a detectable allometric component, as indicated by a significant association with SL (Pillai’s trace = 0.924, F_(16, 10)_ = 7.629, *p* = 0.001), whereas the group × SL interaction was not significant (Pillai’s trace = 2.369, F_(64, 52)_ = 1.181, *p* = 0.269). The exploratory PCA showed separation of *Hypostomus* sp. G1 and *Hypostomus* sp. G2 (hereafter G1 and G2) from *H. niceforoi* and H*. robinii*, but overlap among G1, G2, and *H. rhantos*; PC1 and PC2 explained 30.57% and 14.99% of total variance, respectively (Fig 6A). The corresponding PCA-LDA retained 16 principal components accounting for 95.90% of cumulative variance, with LD1 and LD2 explaining 61.97% and 22.47% of discriminant variation, respectively (Fig 6B). The first discriminant axis separated G1 and G2 from *H. robinii*, whereas the second separated G1 from G2 and *H. niceforoi*. Morphotype G2 overlapped extensively with *H. rhantos* and occupied a position closer to *H. niceforoi* in discriminant space.

**Fig 6.**
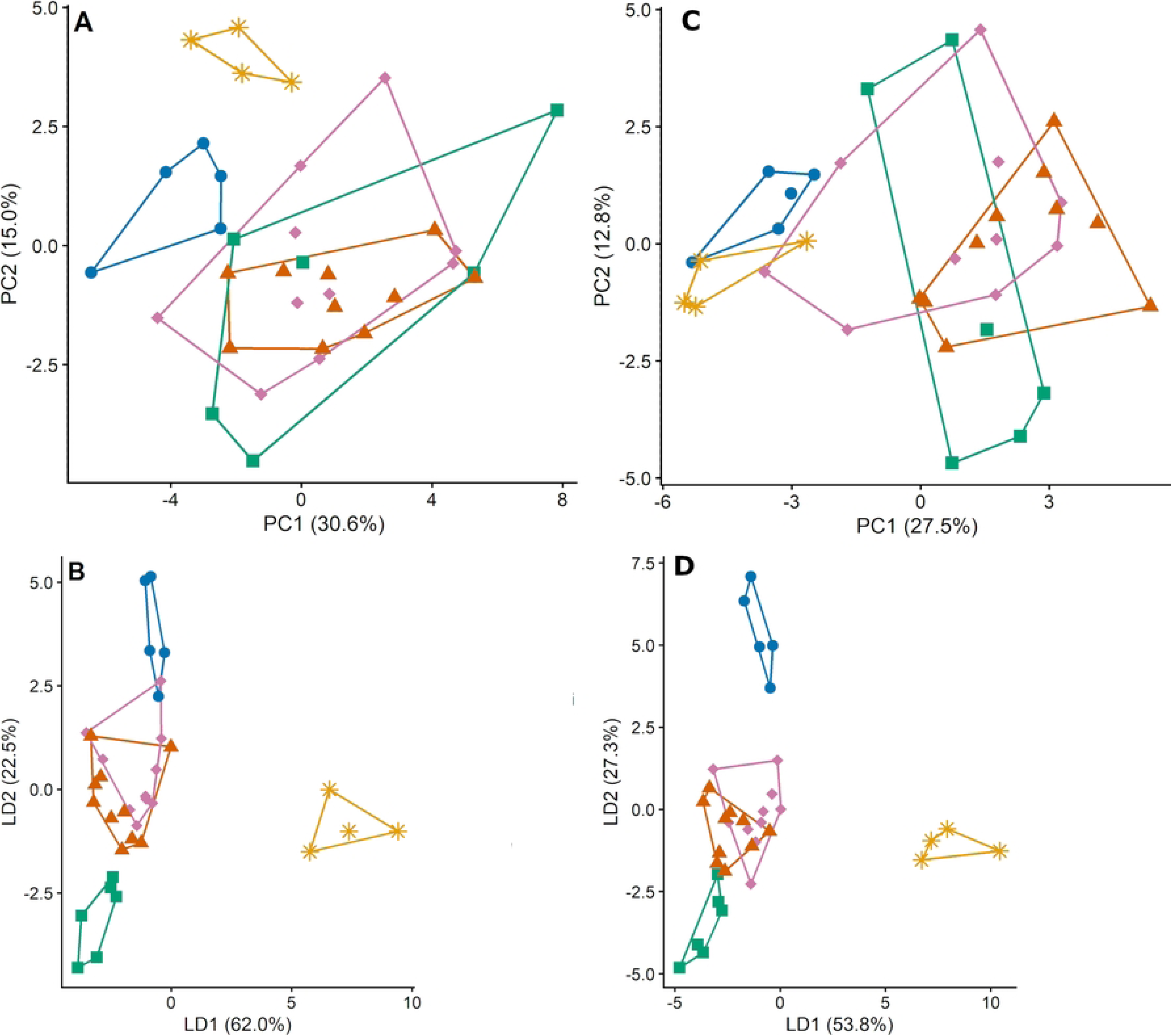
Morphometric variation among *Hypostomus niceforoi*, *H. rhantos*, *H.* sp. G1, *H.* sp. G2, and topotypic *H. robinii*, based on body measurements before and after allometric correction. Exploratory PCA (A) and PCA-LDA (B) used ln(%SL) body measurements without allometric correction. PCA (C) and PCA-LDA (D) used 33 allometrically corrected body variables. The PCA-LDA models retained 16 principal components in B (95.90% of cumulative variance) and 17 in D (95.79%). Symbols indicate *Hypostomus niceforoi* (blue circles, n = 5), *H. rhantos* (orange triangles, n = 10), G1 (green squares, n = 6), G2 (pink diamonds, n = 10), and topotypic *H. robinii* (yellow asterisks, n = 4). Lines outline the convex hulls.

After allometric correction, no residual association with SL was detected (Pillai’s trace = 0.018, F_(17, 9)_ = 0.010, *p* = 1.000), and the group × SL interaction remained nonsignificant (Pillai’s trace = 2.533, F_(68, 48)_ = 1.218, *p* = 0.237). The corrected PCA showed separation of G1 from *H. robinii* and *H. niceforoi* along PC1, but overlap between G1 and *H. rhantos*, with G2 broadly distributed across the space occupied by the other groups. The first two principal components explained 27.47% and 12.84% of total variance, respectively (Fig 6C). The corresponding PCA-LDA retained 17 principal components accounting for 95.79% of cumulative variance, with LD1 and LD2 explaining 55.28% and 25.30% of discriminant variation, respectively (Fig 6D). The first discriminant axis separated G1 and G2 from *H. robinii*, whereas the second separated G1 from G2 and *H. niceforoi*. Morphotype G2 also showed separation from *H. niceforoi* but retained extensive overlap with *H. rhantos*.

In the broad five-group comparison, exploratory analyses of ln(%HL) head measurements retained a detectable allometric component, as indicated by a significant association with HL (Pillai’s trace = 0.908, F_(15, 11)_ = 7.255, *p* = 0.001), whereas the group × HL interaction was not significant (Pillai’s trace = 2.081, F_(60, 56)_ = 1.012, *p* = 0.483). The exploratory PCA showed overlap among groups along PC1, while PC2 separated G1 from *H. robinii* and partially from *H. niceforoi*, but showed overlap with G2 and *H. rhantos*. Morphotype G2 was broadly distributed across the space occupied by the compared groups. The first two principal components explained 31.05% and 21.61% of total variance, respectively (Fig 7A). The corresponding LDA yielded LD1 and LD2 accounting for 54.19% and 24.95% of discriminant variation, respectively (Fig 7B). The first discriminant axis separated G1 from *H. robinii*, *H. niceforoi*, and G2, with G2 overlapping with *H. rhantos*; the second also separated G1 from *H. niceforoi* and *H. robinii*.

**Fig 7.**
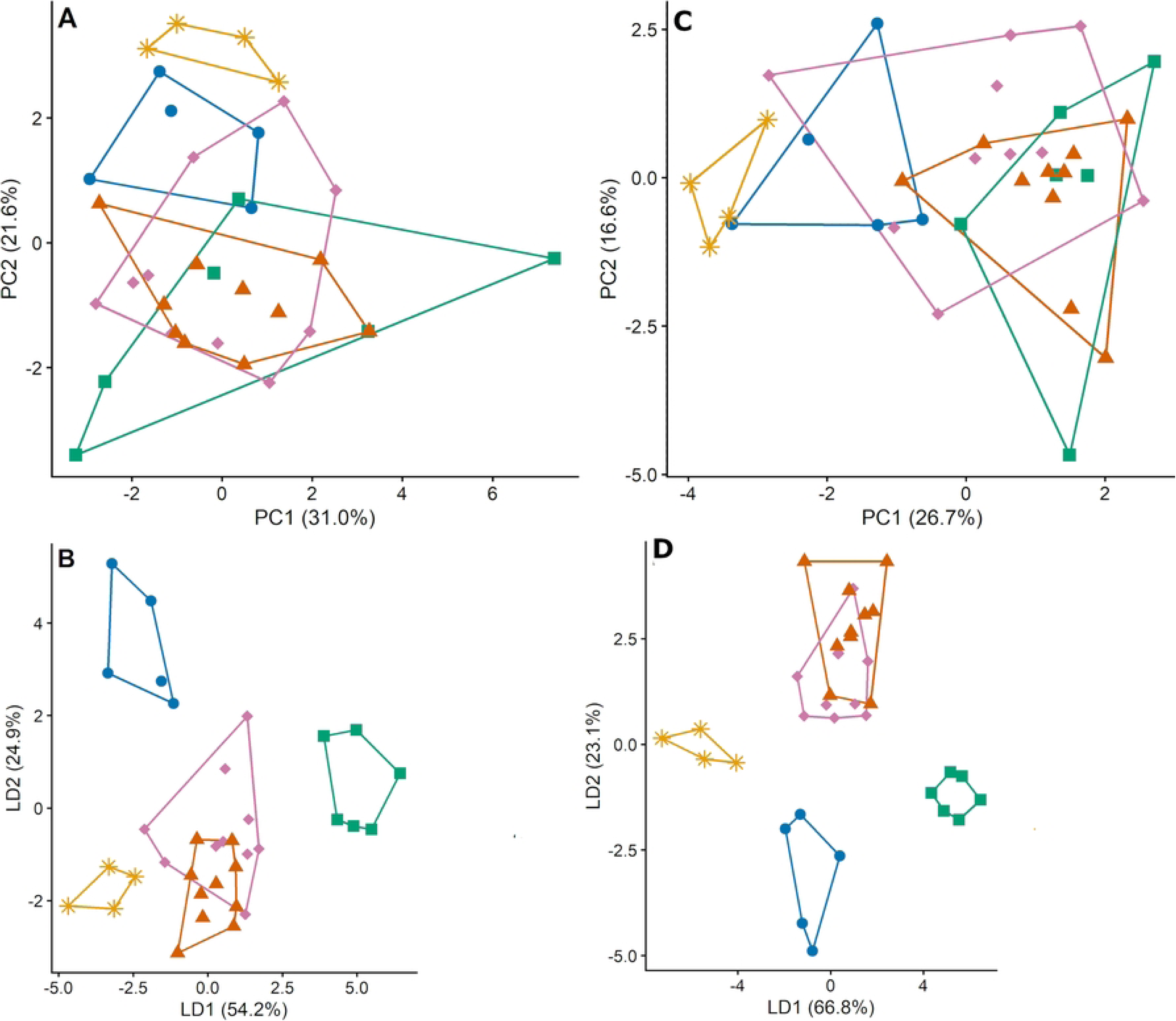
Morphometric variation among *Hypostomus niceforoi*, *H. rhantos*, *H.* sp. G1, *H.* sp. G2, and topotypic *H. robinii*, based on head measurements before and after allometric correction. Exploratory PCA (A) and LDA (B) used 15 ln(%HL) head variables without allometric correction. PCA (C) and LDA (D) used 14 allometrically corrected head variables retained after Holm-adjusted tests of slope homogeneity. Symbols indicate *Hypostomus niceforoi* (blue circles, n = 5), *H. rhantos* (orange triangles, n = 10), G1 (green squares, n = 6), G2 (pink diamonds, n = 10), and topotypic *H. robinii* (yellow asterisks, n = 4). Lines outline the convex hulls.

Predorsal length was excluded because it showed significant heterogeneity of allometric slopes among groups after Holm adjustment. After allometric correction of the remaining 14 variables, no residual association with HL was detected (Pillai’s trace = 0.029, F_(14, 12)_ = 0.026, *p* = 1.000), and the group × HL interaction remained nonsignificant (Pillai’s trace = 1.871, F_(56, 60)_ = 0.942, *p* = 0.588). The corrected PCA showed separation of G1 from *H. robinii* and *H. niceforoi* along PC1, but overlap between G1 and *H. rhantos*, while G2 remained broadly distributed across groups. The first two principal components explained 26.7% and 16.6% of total variance, respectively (Fig 7C). The corresponding LDA yielded LD1 and LD2 accounting for 66.8% and 23.1% of discriminant variation, respectively (Fig 7D). The first discriminant axis separated G1 from *H. robinii*, *H. niceforoi*, and the overlapping G2 and *H. rhantos* groups, whereas the second separated G2 from G1, *H. niceforoi*, and *H. robinii*. Leave-one-out cross-validation with the allometric correction held fixed correctly classified 26 of 35 specimens (overall accuracy = 74.29%; balanced accuracy = 77.00%; Cohen’s κ = 0.669), indicating that separation in the fitted discriminant space coexisted with individual classification errors.

### Focused comparison of G1, G2, and *Hypostomus robinii*

In the focal SL-based comparison of G1, G2, and topotypic *H. robinii*, all 33 body variables were retained because none showed significant heterogeneity of allometric slope after Holm adjustment. Because the corrected body matrix exceeded the residual rank available for direct multivariate testing, residual size effects were evaluated using the first 12 principal components, which explained 95.49% of cumulative variance. No overall residual association with SL was detected (Pillai’s trace = 0.022, F_(12, 3)_ = 0.006, *P* = 1.000), but the group × SL interaction was significant (Pillai’s trace = 1.856, F_(24, 8)_ = 4.296, *P* = 0.019), indicating that residual multivariate relationships with body size differed among groups. The corrected PCA separated G1 from *H. robinii* along PC1, with G2 distributed between them and overlap among groups along PC2; these axes explained 26.16% and 15.66% of total variance, respectively (Fig 8A). The corresponding PCA-LDA, fitted to the 12 retained components, separated G2 from G1 and *H. robinii* along the first discriminant axis and G1 from G2 and *H. robinii* along the second (Fig 8B).

**Fig 8.**
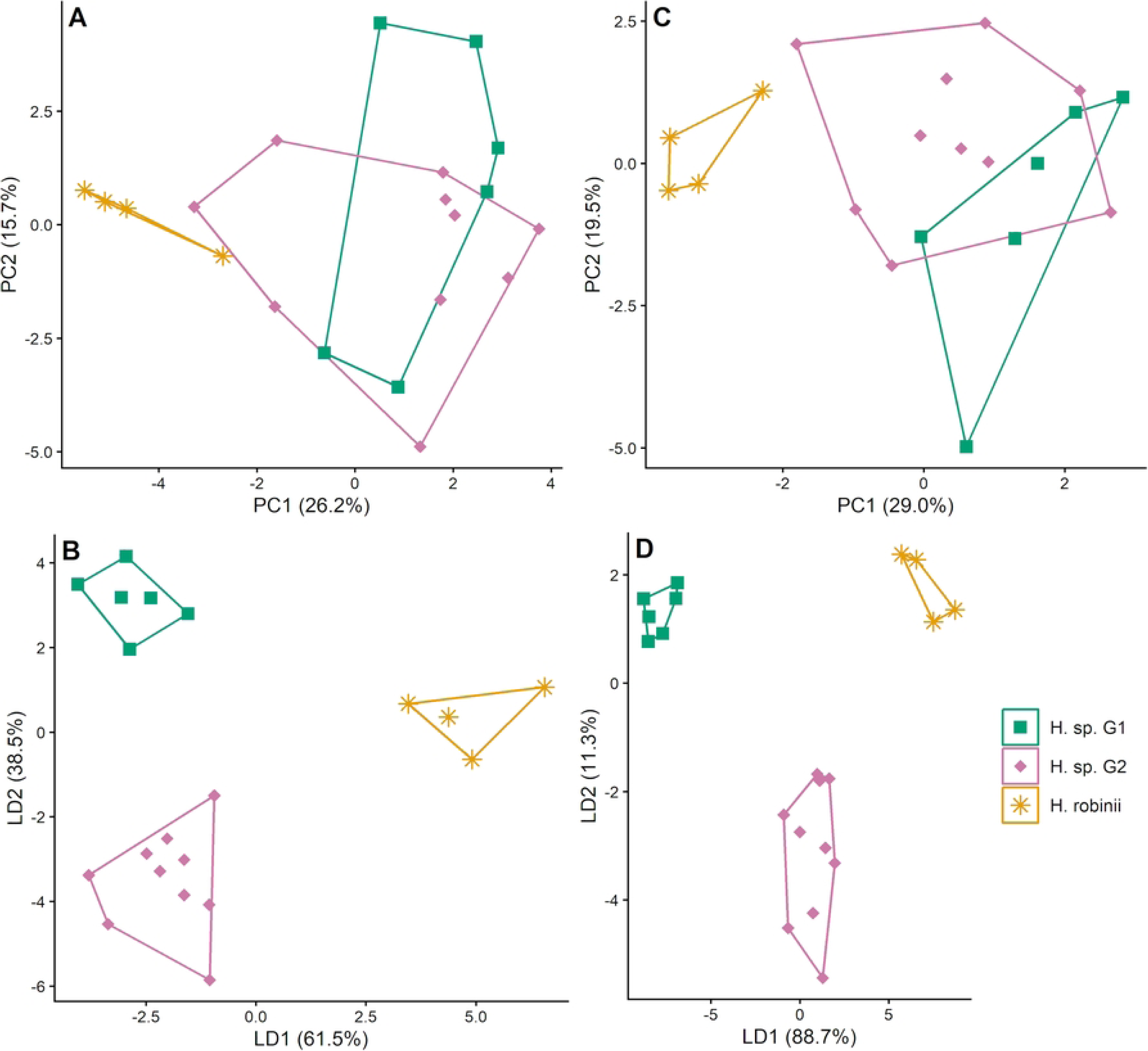
Morphometric variation among *Hypostomus* sp. G1, *H.* sp. G2, and topotypic *H. robinii*, based on allometrically corrected body and head measurements. PCA (A, PC1: 26.16%, PC2: 15.66%) and PCA-LDA (B, 12 PCs retained: 95.49%) used 33 allometrically corrected body variables. PCA (C, PC1: 28.95%, PC2: 19.48%) and LDA (D, LD1: 88.69%, LD2: 11.31%) used 14 allometrically corrected head variables from the same specimens. Percentages refer to total variance explained by the PCs and to discriminant variation accounted for by the LDs. Symbols indicate G1 (green squares, n = 6), G2 (pink diamonds, n = 10), and topotypic *H. robinii* (yellow asterisks, n = 4). Lines outline the convex hulls.

Fully nested leave-one-out cross-validation, with allometric correction and model fitting repeated within each training partition, correctly classified 12 of 20 specimens (overall accuracy = 60.0%; balanced accuracy = 64.44%; Cohen’s κ = 0.355). Recall was 33.33% for G1, 60.0% for G2, and 100% for *H. robinii*, indicating limited predictive discrimination, particularly for G1, despite separation in the fitted discriminant space.

In the focal HL-based comparison, the corrected head matrix comprised 14 variables after exclusion of predorsal length, which showed significant heterogeneity of allometric slopes among groups after Holm adjustment. No residual association with HL was detected (Pillai’s trace = 0.010, F_(8, 7)_ = 0.009, *p* = 1.000), and the group × HL interaction was not significant (Pillai’s trace = 1.146, F_(16, 16)_ = 1.341, *p* = 0.282). The corrected PCA separated G1 and G2 from *H. robinii* along PC1, with overlap between the morphotypes, while PC2 showed a tendency toward G1–G2 separation. These axes explained 29.0% and 19.5% of total variance, respectively (Fig 8C). The corresponding LDA separated G1 and G2 from *H. robinii* along LD1 and G2 from G1 and *H. robinii* along LD2; these axes accounted for 88.7% and 11.3% of discriminant variation, respectively (Fig 8D).

Despite separation in the fitted discriminant space, fully nested LOOCV, with allometric correction and model fitting repeated within each training partition, correctly classified only nine of 20 specimens (overall accuracy = 45.0%; balanced accuracy = 41.67%; Cohen’s κ = 0.134). Correct classifications comprised three of the six G1 specimens (recall = 50.0%), five of the ten G2 specimens (50.0%), and one of the four topotypic *H. robinii* specimens (25.0%).

Univariate comparisons of the 14 corrected head variables detected differences among the three groups in six variables before multiplicity adjustment, four of which remained significant after Holm correction. Snout length and snout–naris length were smaller in G1 than in G2 and *H. robinii*, with no significant differences between G2 and *H. robinii*. Mouth width and dentary length were greater in *H. robinii* than in either morphotype (Table 2).

**TABLE 2.** Head variables showing significant overall differences among *Hypostomus* sp. G1, *Hypostomus* sp. G2, and topotypic specimens of *H. robinii* after Holm correction across 14 univariate tests. Allometrically corrected measurements are expressed in millimeters (mm) and presented as mean (minimum–maximum). The correction used HL as the size variable and the mean HL of the 20-specimen focal reference dataset as the reference length (HL₀). Pairwise comparisons are shown only when both the global test and the corresponding post hoc contrast were significant.

| **Variable/Mean (Range)** | **Snout length** | **Snout–naris length** | **Mouth width** | **Dentary length** |
|---|---|---|---|---|
| *H.* sp. G1 | 24.07 (23.60–24.42) | 15.06 (14.29–15.75) | 17.77 (15.29–21.26) | 5.14 (3.91–6.26) |
| *H.* sp. G2 | 24.78 (24.01–25.89) | 15.94 (15.39–16.73) | 19.25 (16.31–23.69) | 5.48 (4.24–7.96) |
| *H. robinii* (*Hr*) | 25.29 (25.20–25.43) | 16.31 (15.99–16.68) | 23.85 (21.93–24.57) | 8.37 (7.39–9.39) |
| ANOVA | Welch, F<sub>(2, 10.16)</sub> = 42.61 | F<sub>(2, 17)</sub> = 9.61 | F<sub>(2, 17)</sub> = 9.60 | F<sub>(2, 17)</sub> = 13.77 |
| p | 0.00001 | 0.00161 | 0.00162 | 0.00028 |
| Adjusted p (Holm) | 0.00016 | 0.01935 | 0.01935 | 0.00362 |
| Significant pairwise comparisons | G1 < G2; G1 < *Hr* | G1 < G2; G1 < *Hr* | G1 < *Hr*; G2 < *H r* | G1 < *Hr*; G2 < *Hr* |

### External evaluation with Magdalena basin specimens

External evaluation comprised nine Magdalena specimens molecularly assigned to G1 (n = 7) or G2 (n = 2). Seven of the nine specimens exceeded the maximum standard length (SL) of the focal reference sample, and seven exceeded its maximum head length (HL). The LDA model trained exclusively on reference specimens correctly classified two of the nine Magdalena specimens (overall accuracy = 22.22%; balanced accuracy = 14.29%). Two of the seven G1 specimens were correctly classified (recall = 28.57%), whereas neither of the two G2 specimens was correctly classified (recall = 0%). Among the seven specimens molecularly assigned to G1, two were classified morphometrically as G1, three as G2, and two as *H. robinii*. Both G2 specimens were classified as G1 (S3 Table).

### Integrative evaluation of the examined entities

Integration of mitochondrial, morphological, and morphometric evidence revealed different levels of support among the entities examined (Tables 3 and 4).

**TABLE 3.** Molecular delimitation results, phenotypic congruence, and taxonomic assignment support for nominal *Hypostomus* species.

| **Species** | **N** (*cox1*/D-loop) | *cox1* MOTUs |  |  | D-loop MOTUs |  |  | **Phenotypic congruence** | **Taxonomic reference and materials** | **Taxonomic assignment support** |
|---|---|---|---|---|---|---|---|---|---|---|
|  |  | **ASAP** | **bPTP** | **GMYC** | **ASAP** | **bPTP** | **GMYC** |  |  |  |
| *H. holostictus* | 4/4 | 1 | 1 | 1 | 1 | 1 | 1 | C | [16]; photographs of the holotype BMNH 1913.10.1.57 | **H** |
| *H. hondae* | 12/11 | 1 | 1 | 1 | 5 | 5 | 5 | C | [17] | **H** |
| *H. niceforoi* | 4/4 | 1 | 1 | 1 | 2 | 2 | 2 | C | [18]; ANSP 70511, holotype | **H** |
| *H. plecostomoides* | 4/4 | 1 | 1 | 1 | 3 | 3 | 3 | C | [17] | **H** |
| *H. oculeus* | 3/2 | 1 | 1 | 1 | 2 | 1 | 1 | C | [17]; ANSP 70518, holotype | **H** |
| *H. weberi* | 2/4 | 1 | 1 | 1 | 1 | 1 | 2 | C | [20]; examined reference specimens | **H** |
| *H. carinatus* | 1/1 | 1 | 1 | 1 | 1 | 1 | 1 | C | [19] | **M** |
| *H. hemicochliodon* | 1/0 | 1 | 1 | 1* | — | — | — | C | [17] | **M** |
| *H. sculpodon* | 1/0 | 1 | 1 | 1 | — | — | — | C | [17] | **M** |
| *H. wilsoni* | 2/0 | 1 | 1 | 1 | — | — | — | C** | [21]; photographs of the holotype FMNH 58550; holotype of *H. aspidolepis* BMNH 1866.11.5.1; [22] | **P** |

N (cox1/D-loop), number of individuals sequenced for each marker; MOTU, molecular operational taxonomic unit; C, congruence with diagnostic phenotypic characters; H, high support; M, moderate support; P, provisional support. Taxonomic assignment support reflects confidence in specimen identification based on diagnostic phenotypic congruence, taxonomic reference material, sample representation, and mitochondrial partition compatibility, following the operational criteria detailed in S4 Table. These categories do not imply evolutionary independence or formal species recognition. An em dash (—) indicates unavailable data. \*Discordant MOTU composition relative to the other delimitation methods. \*\*Sequenced individuals lacking associated voucher specimens.

**TABLE 4.**
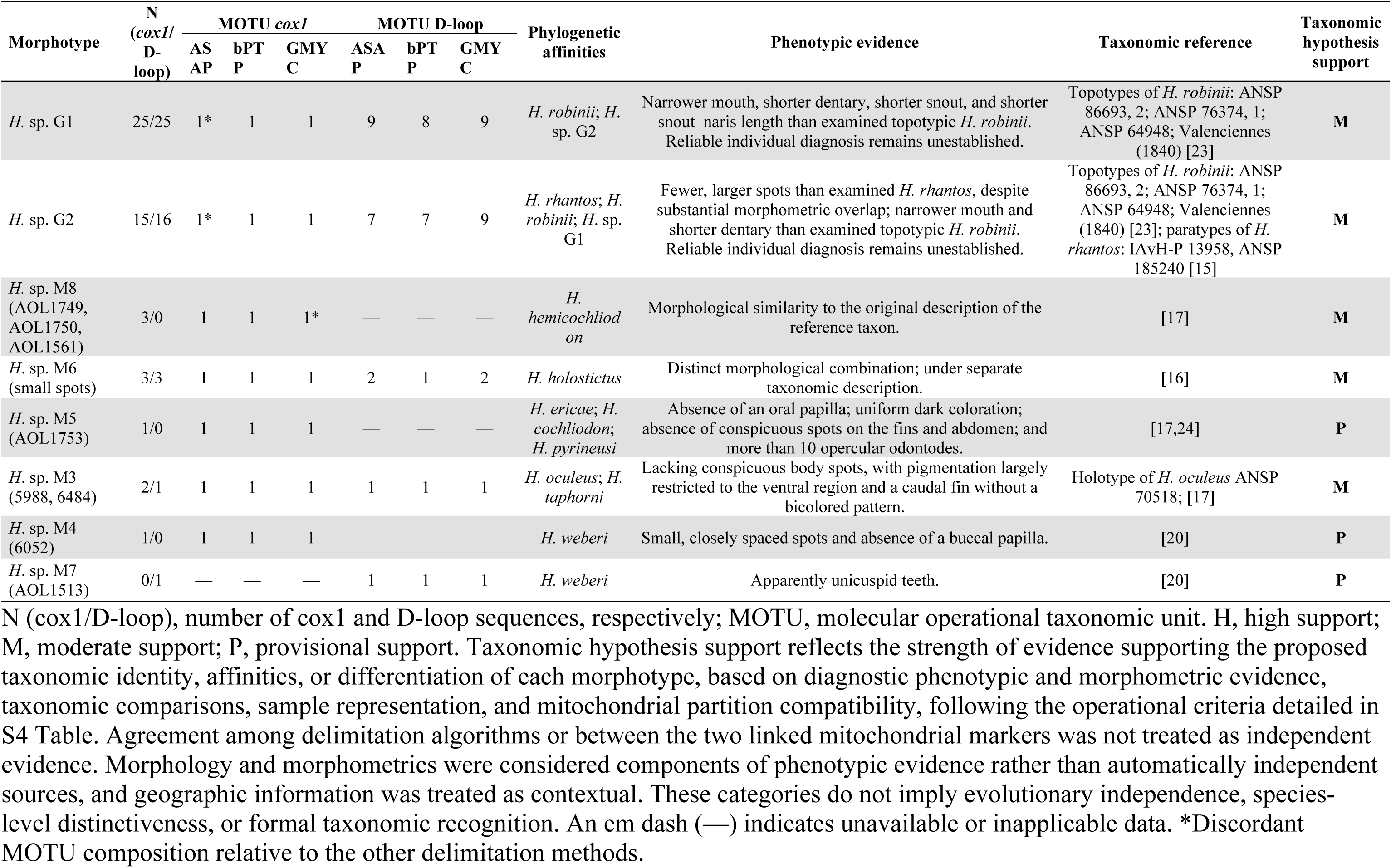
Summary of molecular, phenotypic, and taxonomic evidence for the *Hypostomus* morphotypes evaluated.

| Morphotype | N (*cox1*/D-loop) | MOTU *cox1* |  |  | MOTU D-loop |  |  | Phylogenetic affinities | Phenotypic evidence | Taxonomic reference | Taxonomic hypothesis support |
|---|---|---|---|---|---|---|---|---|---|---|---|
|  |  | ASAP | bPTP | GMYC | ASAP | bPTP | GMYC |  |  |  |  |
| *H.* sp. G1 | 25/25 | 1* | 1 | 1 | 9 | 8 | 9 | *H. robinii*; *H.* sp. G2 | Narrower mouth, shorter dentary, shorter snout, and shorter snout–naris length than examined topotypic *H. robinii*. Reliable individual diagnosis remains unestablished. | Topotypes of *H. robinii*: ANSP 86693, 2; ANSP 76374, 1; ANSP 64948; Valenciennes (1840) [23] | **M** |
| *H.* sp. G2 | 15/16 | 1* | 1 | 1 | 7 | 7 | 9 | *H. rhantos*; *H. robinii*; *H.* sp. G1 | Fewer, larger spots than examined *H. rhantos*, despite substantial morphometric overlap; narrower mouth and shorter dentary than examined topotypic *H. robinii*. Reliable individual diagnosis remains unestablished. | Topotypes of *H. robinii*: ANSP 86693, 2; ANSP 76374, 1; ANSP 64948; Valenciennes (1840) [23]; paratypes of *H. rhantos*: IAvH-P 13958, ANSP 185240 [15] | **M** |
| *H.* sp. M8 (AOL1749, AOL1750, AOL1561) | 3/0 | 1 | 1 | 1* | — | — | — | *H. hemicochliodon* | Morphological similarity to the original description of the reference taxon. | [17] | **M** |
| *H.* sp. M6 (small spots) | 3/3 | 1 | 1 | 1 | 2 | 1 | 2 | *H. holostictus* | Distinct morphological combination; under separate taxonomic description. | [16] | **M** |
| *H.* sp. M5 (AOL1753) | 1/0 | 1 | 1 | 1 | — | — | — | *H. ericae*; *H. cochliodon*; *H. pyrineusi* | Absence of an oral papilla; uniform dark coloration; absence of conspicuous spots on the fins and abdomen; and more than 10 opercular odontodes. | [17,24] | **P** |
| *H.* sp. M3 (5988, 6484) | 2/1 | 1 | 1 | 1 | 1 | 1 | 1 | *H. oculeus*; *H. taphorni* | Lacking conspicuous body spots, with pigmentation largely restricted to the ventral region and a caudal fin without a bicolored pattern. | Holotype of *H. oculeus* ANSP 70518; [17] | **M** |
| *H.* sp. M4 (6052) | 1/0 | 1 | 1 | 1 | — | — | — | *H. weberi* | Small, closely spaced spots and absence of a buccal papilla. | [20] | **P** |
| *H.* sp. M7 (AOL1513) | 0/1 | — | — | — | 1 | 1 | 1 | *H. weberi* | Apparently unicuspid teeth. | [20] | **P** |

N (cox1/D-loop), number of cox1 and D-loop sequences, respectively; MOTU, molecular operational taxonomic unit. H, high support; M, moderate support; P, provisional support. Taxonomic hypothesis support reflects the strength of evidence supporting the proposed taxonomic identity, affinities, or differentiation of each morphotype, based on diagnostic phenotypic and morphometric evidence, taxonomic comparisons, sample representation, and mitochondrial partition compatibility, following the operational criteria detailed in S4 Table. Agreement among delimitation algorithms or between the two linked mitochondrial markers was not treated as independent evidence. Morphology and morphometrics were considered components of phenotypic evidence rather than automatically independent sources, and geographic information was treated as contextual. These categories do not imply evolutionary independence, species-level distinctiveness, or formal taxonomic recognition. An em dash (—) indicates unavailable or inapplicable data. \*Discordant MOTU composition relative to the other delimitation methods.

Among nominal species, *H. holostictus*, *H. hondae*, *H. niceforoi*, *H. plecostomoides*, *H. oculeus*, and *H. weberi* received high taxonomic-assignment support under the operational criteria in S4 Table. Direct comparison with holotype material provided strong taxonomic corroboration for *H. niceforoi* and *H. oculeus*, supporting the latter assignment despite limited sampling. For *H. weberi*, diagnostic phenotypic congruence, comparative reference material, and representation by four distinct individuals supported the identification. Mitochondrial subdivision was not considered evidence against these assignments in the absence of documented conflicts with diagnostic phenotypic or taxonomic evidence. *Hypostomus carinatus* received moderate support because it was represented by a single individual; *H. hemicochliodon*, and *H. sculpodon* were additionally limited by the absence of D-loop data. The assignment of *H. wilsoni* remained provisional because the absence of vouchers directly associated with the sequenced tissues limited independent verification.

Among the evaluated morphotypes, G1 and G2 had the broadest documented evidence base but received moderate taxonomic-hypothesis support. Both were characterized using mitochondrial markers and comparisons with relevant taxonomic material. Phenotypic evidence included differences in selected morphometric characters relative to examined topotypic *H. robinii* and qualitative pigmentation differences between G2 and *H. rhantos*. However, these contrasts did not establish reliable individual diagnoses. In the external evaluation of nine Magdalena specimens, the reference-trained LDA model achieved an overall accuracy of 22.22% and a balanced accuracy of 14.29%, with errors involving confusion between G1 and G2 and assignment of some G1 specimens to *H. robinii*. This limited external classification performance, together with incomplete diagnostic resolution and unresolved specific identities, precluded high support. *Hypostomus* sp. M6 and *Hypostomus* sp. M3 also received moderate support. M6 showed a distinctive morphological combination and was represented by multiple specimens, whereas M3 combined phenotypic differences from related taxa with taxonomic comparisons and mitochondrial characterization, although its limited sampling constrained assessment of diagnostic stability. *Hypostomus* sp. M4, sp. M5, and sp. M7 remained provisional because their proposed differences were supported by one or very few specimens. M8 received moderate support for identification as *H. hemicochliodon* and was not treated as an unresolved candidate lineage because the examined specimens were morphologically congruent with that species.

## Discussion

Integrating mitochondrial, qualitative morphological, and morphometric evidence revealed different levels of correspondence between Colombian *Hypostomus* entities and currently recognized species. Several nominal identifications were supported by diagnostic morphology and reference comparisons, whereas other entities remained unresolved despite detectable mitochondrial or phenotypic differentiation. The principal contrast was between relatively coarse *cox1* partitions and substantially finer D-loop subdivision. *Hypostomus* sp. G1 and G2 further illustrate the difficulty of translating mitochondrial affinities and phenotypic contrasts with reference material into definitive taxonomic assignments. These findings support an integrative assessment in which mitochondrial partitions generate hypotheses, while their taxonomic interpretation depends on the nature, independence, and geographic coverage of the available evidence [10].

### Mitochondrial partitioning and its taxonomic interpretation

The observed variation is consistent with the difficulty of delimiting species in *Hypostomus* using mitochondrial divergence alone. A continental multilocus analysis found that morphologically recognized sister species often showed *cox1* divergence below the conventional 2% barcode threshold [2]. Conversely, integrative revision of the *H. pusarum* complex resolved several nominal taxa as synonyms while recognizing an additional species [64]. Thus, integrative assessment can reveal overlooked diversity or reduce taxonomic overestimation; neither low mitochondrial divergence nor a molecular discontinuity determines the taxonomic outcome in advance.

D-loop recovered substantially more MOTUs than *cox1*, indicating finer partitioning of the sampled mitochondrial variation. This pattern was also observed among specimens represented by both markers, indicating that unequal specimen representation alone cannot explain the difference. Differences in sequence variability may contribute to the contrasting partitions recovered from the two markers, although their relative substitution rates were not estimated. Teleost mitochondrial control regions contain both conserved and highly variable components, whose organization and variability differ among taxa and sequence segments [65]. Consequently, D-loop partitions may reflect population structure, phylogeographic history, and demographic processes in addition to taxonomic divergence [31,66]. More broadly, mitochondrial genealogies may differ from species histories because of population structure, introgression, retention of ancestral polymorphism, and other evolutionary processes [12,67].

Subdivision within *H. niceforoi*, *H. hondae*, and *H. plecostomoides* illustrates the distinction between mitochondrial structure and nominal identification. Each was recovered as a single *cox1* unit and showed morphological correspondence with its assigned name, whereas D-loop recovered multiple units. These additional partitions corresponded to geographic sampling: Orteguaza and Cohembi localities in *H. niceforoi*; Atrato, middle and lower Magdalena, lower Cauca, and Catatumbo localities in *H. hondae*; and distinct Orinoco localities in *H. plecostomoides*. This correspondence suggests that the finer partitioning recovered by D-loop reflects geographically structured mitochondrial variation within the currently identified species. However, the absence of documented phenotypic differentiation does not exclude unrecognized diversity, particularly where sampling is limited. This possibility is illustrated by cytogenetic evidence of cryptic diversity among morphologically similar populations assigned to *Hypostomus tietensis* [68]. The appropriate interpretation is therefore to retain the nominal identifications while leaving the taxonomic significance of the internal partitions unresolved.

Beyond differences between mitochondrial markers, variation among delimitation methods also affects the interpretation of the recovered MOTUs. ASAP partitions pairwise genetic distances, PTP models branching patterns in substitution units, and GMYC identifies a transition between branching processes on an ultrametric genealogy [6–8]. Their disagreement in the G1–G2, *H. aspidolepis*–*H. wilsoni*, and *H. hemicochliodon*–M8 comparisons identifies partitions sensitive to analytical assumptions. Agreement indicates stability across the applied methods, but does not supply independent biological replication. Likewise, *cox1* and D-loop are linked mitochondrial regions, so concordance between them cannot be treated as corroboration by independently inherited loci [12,67].

GMYC likelihood and threshold estimates describe the fitted genealogies; they do not validate every recovered entity. The performance of single-locus delimitation depends on sampling and evolutionary conditions, including population size relative to divergence time [6,69]. These limitations do not establish that a particular partition is incorrect, but restrict the conclusions that follow from it. Similarly, threshold estimates from separately scaled genealogies cannot be compared directly as divergence dates. Thus, the recovered mitochondrial partitions should be regarded as hypotheses for further population-level and integrative taxonomic investigation rather than direct estimates of species richness.

### Nominal identifications and unresolved morphotypes

Direct comparisons with taxonomic reference specimens provide an essential link between lineages and nominal species assignments [70]. Examination of the *H. niceforoi* and *H. oculeus* holotypes strengthened the corresponding identifications, while diagnostic morphology supported assignments to *H. hondae*, *H. plecostomoides*, *H. holostictus*, and *H. weberi*. For *H. weberi*, reference comparisons and representation by four distinct individuals provided a stronger basis than mitochondrial similarity alone. Under the operational rubric used here, high taxonomic-assignment support refers to identification of the examined material; it does not constitute a comprehensive test of species boundaries across the entire distribution of each species.

Accordingly, D-loop subdivision in *H. niceforoi*, *H. hondae*, and *H. plecostomoides*, and method-dependent subdivision in *H. oculeus* and *H. weberi*, do not by themselves invalidate those assignments. Conversely, the absence of subdivision in the sampled *H. holostictus* cannot establish genetic homogeneity throughout its range. Support for *H. carinatus*, *H. hemicochliodon*, and *H. sculpodon* remains more limited because specimen representation or marker coverage is restricted. For *H. wilsoni*, the lack of vouchers directly associated with sequenced tissues presents a different problem: independently examined specimens and type photographs cannot verify the phenotype–sequence association of those particular samples [71]. The relationship between the sequences assigned to *H. wilsoni* and *H. aspidolepis* therefore remains unresolved.

Among the non-nominal entities, combinations of pigmentation, oral morphology, dentition, and odontodes provide testable hypotheses beyond mitochondrial affinity. For M3, the reported contrasts with *H. oculeus* and *H. taphorni*, including pigmentation and caudal coloration, justify continued comparative evaluation. M4 and M7 also warrant examination relative to *H. weberi*. Its original description emphasizes conspicuous, widely separated rounded spots, bicuspid teeth, and a buccal papilla [20]. The smaller, more closely spaced spots and absence of a papilla reported for M4, and the apparently unicuspid teeth reported for M7, identify potentially informative differences. However, dental interpretation must distinguish absence of a cusp from reduction, fusion, or wear. Substantial intraspecific variation in tooth morphology, buccal-papilla condition, and pigmentation has been documented in *Hypostomus khimaera*, underscoring the need for caution when interpreting these characters [72]. The limited material available here prevents establishing the stability of these differences or of the combination of coloration, oral-papilla condition, and opercular odontodes described for M5. Such comparisons should encompass the variation documented within the *H. cochliodon* group [17].

M6 combines representation by multiple specimens, recovery as a *cox1* unit, and reported dental differentiation from the examined *H. holostictus*. These features support its treatment as a priority taxonomic hypothesis. Nevertheless, the diagnostic interpretation of its bicuspid teeth requires confirmation across specimens and comparison with variation in the reference material. Its method-dependent D-loop subdivision does not independently justify additional entities within M6. Likewise, the mitochondrial placement of M6 and *H. holostictus* outside the supergroups used in the comparative framework does not establish a new supergroup; that possibility requires broader taxon sampling and independently inherited loci.

M8 differs from these cases because no phenotypic distinction from *H. hemicochliodon* has been documented. Its separation by ASAP and bPTP, but grouping with that species by GMYC, therefore supports an unresolved mitochondrial partition rather than a separate taxonomic entity on the present evidence. Morphological correspondence supports its current assignment to *H. hemicochliodon*, while leaving possible population differentiation open to investigation. Distinguishing support for a nominal assignment from support for a new taxonomic hypothesis avoids treating every mitochondrial subdivision as an equivalent taxonomic proposition.

### Morphometric evidence in the G1–G2 complex

*Hypostomus* sp. G1 and G2 have the broadest comparative evidence among the unresolved entities, but their specific identities remain uncertain. Their separation by bPTP and GMYC and grouping by ASAP indicate method-dependent *cox1* differentiation. D-loop subdivided each group much more extensively, paralleling the fragmentation observed within well-supported nominal species. These internal partitions cannot therefore be interpreted directly as additional species. Affinities with *H. robinii* and *H. rhantos* provide a comparative framework rather than definitive identifications.

The morphometric analyses add phenotypic evidence, but their interpretation depends on size adjustment and validation design. Expressing measurements as percentages of standard length or head length did not remove all size-related variation, consistent with the distinction between isometric ratios and allometric normalization [49]. Persistence of some contrasts after adjustment suggests that size alone did not explain the original patterns. Nevertheless, the significant group × standard-length interaction in the focal body dataset indicates group-dependent residual size relationships despite the absence of an overall residual association with size. Those contrasts remain conditional on the sampled size ranges and correction procedure. The absence of a detected interaction in the focal head dataset does not establish identical allometric relationships.

A further distinction concerns separation in fitted discriminant space and prediction of withheld specimens. The broad analyses displayed clearer separation in some discriminant plots than in PCA, while G2 retained substantial overlap with *H. rhantos*. In the focal three-group comparisons with topotypic *H. robinii*, fully nested validation yielded balanced accuracies of 64.44% for body measurements and 41.67% for head measurements. Thus, visually clear fitted separation, especially in the head analysis, did not translate into consistently reliable identification. Repeating model-development steps within each cross-validation training partition is essential to avoid information leakage and overly optimistic estimates of predictive performance [73]. The broad head analysis with allometric correction fixed during LOOCV consequently provides a conditional assessment and should not be equated with the fully nested focal evaluations.

Comparisons with *H. robinii* nevertheless identified informative contrasts. *Hypostomus* sp. G1 and G2 had smaller corrected mouth width and dentary length than the examined topotypes, and G1 also had shorter snout and snout–naris lengths. These findings argue against assigning all examined specimens to *H. robinii* solely on general resemblance or mitochondrial affinity. However, the reference sample comprised only four topotypes, and their classification depended strongly on the measurement dataset: all four were correctly classified using body measurements, compared with one using head measurements. The results therefore support differences from the examined reference material more strongly than a stable diagnosis relative to the full variation of *H. robinii*.

For *H. rhantos*, morphometric similarity must be considered alongside pigmentation. *Hypostomus* sp. G2 overlapped extensively with that species in the measured dimensions, whereas the apparent separation of *Hypostomus* sp. G1 depended on the analysis. Nevertheless, the examined G2 specimens had fewer and larger spots than the compared *H. rhantos*. The original diagnosis of *H. rhantos* emphasizes dense, very small spots and uses size-dependent spot counts [15]. Pigmentation thus contributes evidence beyond the linear measurements, but its diagnostic value requires standardized comparison across body regions, sizes, and preservation conditions. Neither the observed morphometric overlap nor the pigmentation difference resolves whether *Hypostomus* sp. G2 is conspecific with *H. rhantos*.

The three-group comparisons with topotypic *H. robinii* also identified differences in corrected snout length and snout–naris length between G1 and G2. These contrasts provide candidate characters for further evaluation, but their interpretation remains conditional on the sampled specimens and analytical framework. They do not establish diagnostic thresholds or demonstrate reliable individual discrimination between G1 and G2. Moreover, the two groups were represented by different Orinoco localities, preventing the observed contrasts from being attributed exclusively to lineage identity. Replicated sampling across localities, including comparisons within shared geographic settings where possible, is needed to assess the consistency and taxonomic relevance of these differences.

The external evaluation with Magdalena specimens revealed limited diagnostic applicability of the focal three-class head LDA model. Only two of nine specimens were correctly classified, with low recall for both G1 (28.57%) and G2 (0%). Misclassification involved both confusion between G1 and G2 and assignment of two molecularly identified G1 specimens to *H. robinii*. Thus, the morphometric decision boundaries estimated from the focal reference sample did not reliably recover molecular group assignments in the Magdalena specimens. The external evaluation also involved extrapolation beyond the reference size range, as seven of the nine Magdalena specimens exceeded the maximum standard length (SL) and seven exceeded the maximum head length (HL) represented in the focal reference sample. Because the external specimens differed from the reference material in both geographic origin and size coverage, the present design does not allow the potential contributions of these factors to classification performance to be disentangled. The limited external performance restricts the applicability of the model for individual identification beyond the reference sample, but neither establishes conspecificity among the confused groups nor invalidates the phenotypic contrasts observed in the reference material. Broader geographic sampling encompassing a wider range of body sizes, together with validation designs that account for spatial structure, is needed to determine whether these phenotypic contrasts provide consistent diagnostic information across localities [74].

### Geographic context and priorities for integrative assessment

The occurrence of G1 and G2 in both Orinoco and Magdalena, together with the wider regional representation of G1, indicates that their mitochondrial distributions do not map one-to-one onto major drainage boundaries. This pattern motivates investigation of drainage history and connectivity, but does not identify a particular hydrogeological event or distinguish historical connections, retention of ancestral variation, introgression, and recent movement. Recent human-mediated translocation also remains an untested possibility that would require evidence concerning the sampled entities and potential source populations. These alternatives remain hypotheses, not explanations demonstrated by the present mitochondrial and morphometric data.

Future sampling should therefore replicate G1 and G2 across localities and, wherever possible, compare them within shared geographic settings. Such a design would address the present confounding of locality and group, while geographically structured validation would evaluate transfer to unsampled populations. Broader size coverage, standardized preservation and measurement procedures, and additional topotypic *H. robinii* and comparative *H. rhantos* material are needed to determine whether the observed contrasts are stable. Nuclear or genomic data would provide independently inherited evidence for evaluating population differentiation, historical divergence, and the possibility of gene flow.

The study contributes mitochondrial data linked, in most cases, to morphological and geographic information across multiple Colombian hydrographic regions. Its strongest taxonomic conclusions concern supported nominal identifications and documented differences from examined reference material. Under the operational rubric, moderate taxonomic-hypothesis support for G1, G2, M3, and M6 identifies priorities for further evaluation rather than demonstrated new species; M4, M5, and M7 remain more provisional because of restricted representation. Explicit sequence–voucher associations and larger comparative series are essential for testing these hypotheses. Together, these findings characterize mitochondrial variation and clarify the taxonomic relevance of the recovered partitions by distinguishing supported nominal assignments from unresolved morphotype hypotheses and phenotypic contrasts whose diagnostic applicability remains to be established.

## Acknowledgments

We are grateful to the curators, collection managers, staff, and researchers of CIACOL, CIUA, CZUT-IC, IAvH-P, and MPUJ for facilitating access to tissue samples and, in the case of CIUA, for their assistance with the examination of voucher specimens. At CIACOL, we especially thank Astrid Acosta and Juan David Bogotá-Gregory; at CIUA, Luz Fernanda Jiménez and Juan Guillermo Ospina; at CZUT-IC, Francisco A. Villa-Navarro and Harold Flórez-Herrera; at IAvH-P, Ángela Liliana Gutiérrez-Cortés, Yuliana Chala, and Robinson Sánchez; and at MPUJ, Saúl Prada-Pedreros, Javier Farfán, and Santiago Garzón.

## SUPPLEMENTARY INFORMATION

**S1 Table. List of sequences generated in this study and retrieved from GenBank for molecular species delimitation analyses**.

**S2 Table. Raw morphometric data for *Hypostomus* specimens included in comparative analyses.** Specimen and tissue identifiers, taxonomic or morphotype assignments, hydrographic regions, and measurements (mm), organized into standard length (SL)- and head length (HL)-based datasets.

**S3 Table. Individual morphometric predictions for nine Magdalena basin specimens from the three-class head LDA model fitted to the focal reference dataset.** The reference dataset comprised Orinoco G1 (n = 6), Orinoco G2 (n = 10), and topotypic *H. robinii* (n = 4), and the model used the 14 retained allometrically corrected head variables. The Magdalena specimens were not used in allometric preprocessing, variable screening, model fitting, or internal validation. External specimens were transformed using parameters estimated exclusively from the 20-specimen focal reference dataset. Independent molecular assignments to G1 or G2 were incorporated only after morphometric prediction to evaluate agreement. Predictions as *H. robinii* represent assignments within the fitted morphometric model and are not interpreted as taxonomic identifications.

**S4 Table. Operational criteria and evidence for taxonomic assignments of nominal *Hypostomus* species and hypotheses for unidentified morphotypes.** Qualitative support categories—high (H), moderate (M), and provisional (P)—are defined with decision rules and taxon-specific evidence and limitations informing the integrative evaluation.

